# Purification of micrococcal nuclease (MNase) for use in ribosomal profiling of high-salinity extremophiles

**DOI:** 10.1101/2024.10.03.616411

**Authors:** Pavlina Gregorova, Matthew Isada, Jocelyne DiRuggiero, L. Peter Sarin

**Affiliations:** RNAcious Laboratory, Molecular and Integrative Biosciences Research Programme, Faculty of Biological and Environmental Sciences, University of Helsinki, Viikinkaari 9B, 00790 Helsinki, Finland; Department of Biology, Johns Hopkins University, 3400 N. Charles St., 235 Mudd Hall, Baltimore, MD 21218 Baltimore, Maryland, USA

**Keywords:** Micrococcal nuclease, S7, MNase, ribosome profiling, haloarchaea, halophiles, archaea

## Abstract

Nucleases, i.e. enzymes that catalyze the hydrolysis of phosphodiester bonds in nucleic acids, are essential tools in molecular biology and biotechnology. *Staphylococcus aureus nuclease (M*Nase) is particularly interesting due to its thermostability and Ca^2+^-dependence, making it the prime choice for applications where nuclease modulation is critical, such as ribosome profiling in bacteria and halophilic archaea. The latter poses a technical and economical challenge: high salt reaction conditions are essential for maintaining ribosome integrity but negatively impact the MNase activity, necessitating using large amounts of nuclease to achieve efficient cleavage. Here, we set out to generate an optimized production protocol for two forms of MNase — fully processed MNaseA and the 19 amino acid propeptide-containing MNaseB — and to biochemically benchmark them against a commercial nuclease. Our results show that both MNases are highly active in normal reaction conditions, but MNaseA maintains higher enzymatic activity in high salt concentrations than MNaseB. MNaseA also retains >90% of its activity after multiple freeze-thaw cycles when stored at -80 °C in a buffer containing 5% glycerol. Importantly, ribosome profiling experiments in the haloarchaeon *Haloferax volcanii demonstrate*d that MNaseA produces ribosome footprints and hallmarks of active translation highly comparable to those obtained with the commercial nuclease, making it a suitable alternative for high-salt ribosome profiling applications. In conclusion, our method can be easily implemented for efficient MNaseA production, thereby providing access to an effective, robust, and cost-efficient alternative to commercial nucleases, as well as facilitating future translation studies into halophilic organisms.

## INTRODUCTION

Nucleases constitute a diverse family of enzymes with various biochemical and catalytic properties that mediate the degradation of nucleic acids by cleaving the phosphodiester bonds of their backbone (1). Extracellular nucleases, which can be membrane-bound or secreted, are often important virulence factors in bacterial pathogenesis. For example, *Staphylococcus aureus nuclease (N*uc) is required for the evasion of neutrophil extracellular traps and the inhibition of biofilm formation through the cleavage of extracellular DNA (2). Nuc was identified by Cunningham *et al. (3) and ter*med micrococcal nuclease (abbreviated MNase). It is a thermostable, strictly Ca^2+^-dependent phosphodiesterase that hydrolyses single or double-stranded RNA or DNA to produce 3’-phosphomononucleotides (4, 5). The *nuc gene encode*s for a protein with a 60-residue long secretory signaling sequence, which is subsequently removed during secretion. The secreted propeptide form of MNase, known as MNaseB, is further processed by most *S. aureus strains to* yield the mature form, MNaseA, which is 19 residues shorter. Interestingly, both MNaseA and MNaseB are enzymatically active (6). Moreover, *S. aureus expresses a* second endonuclease, Nuc2, which in contrast, is membrane-bound and mainly associated with biofilm regulation (7).

The thermostability and strict Ca^2+^-dependence make MNase an attractive nuclease for research and biotechnological applications. An early use of MNase included the degradation of chromatin to elucidate nucleosome occupancy (8), which has subsequently been combined with high-throughput sequencing to yield various chromatin profiling strategies, such as pinpointing transposase-accessible chromatin sites for epigenetic analyses (9) and antibody-targeted *in situ mapping of* protein-DNA interactions (10). Furthermore, Pelham and Jackson utilized MNase to perform a limited nuclease treatment to remove endogenous messenger RNA (mRNA) from cell-free extracts without damaging other RNA components, such as transfer RNA (tRNA) and ribosomal RNA (rRNA). This is achieved by terminating nuclease activity by adding EGTA, which preferentially chelates Ca^2+^ ions over other divalent cations (11, 12). MNase has also been used to improve downstream processing in recombinant protein expression, whereby a modified *Escherichia coli JM107 strai*n expresses extracellular MNaseA to reduce sample viscosity upon cell lysis by hydrolyzing the host’s chromosomal DNA (13).

The development of ribosome profiling — a method for monitoring translation across the translatome with subcodon resolution — has emphasized the need for a ribonuclease that efficiently hydrolyzes mRNA outside of the translating ribosomes without damaging the ribosomes. The original method by Ingolia *et al. (14) succ*essfully utilized RNase I to generate ribosome protected footprints in *Saccharomyces cerevisiae; RNase I has* since been the nuclease-of-choice for most ribosome profiling applications. However, RNase I has been shown to severely degrade ribosomes in some species and other nucleases, such as MNase, may provide less aggressive alternatives suitable for eukaryotes and bacteria (15). To date, MNase is primarily used in bacterial ribosome profiling studies, where *E. coli RNase I is* incompatible as its nuclease activity is inhibited by *E. coli ribosomes (*16, 17). Contrary to RNase I, MNase also retains some activity in high-salinity conditions, which makes it a suitable choice for studies on halophilic organisms, such as the model archaeon *Haloferax volcanii (18)*.

Although MNase is the preferred choice for prokaryotic ribosome profiling (Ribo-seq), there are some limitations to consider. MNase solutions are available from several commercial vendors, but most products are supplied at a concentration too low to achieve the required activity, especially when working with halophilic organisms. Lyophilized MNase batches provide sufficient activity but cost quickly becomes a limiting factor when multiple samples are processed. An alternative approach is to produce MNase in-house, which presents its challenges, as expressing an active nuclease is toxic to the expression strain and limits protein yields. This can be largely circumvented by introducing a secretory signaling sequence and targeting the recombinant protein for secretion to the periplasm, which furthers host viability and improves enzyme activity (19). OmpA-directed periplasmic targeting has been used to express both MNaseA (20, 21) and MNaseB (20, 22) in various *E. coli strains. Si*milar approaches have been demonstrated for other bacterial hosts. The native form of MNase was successfully expressed and secreted in *Corynebacterium glutamicum using a sta*phylococcal-derived vector (23). However, MNase bears an atypical signaling sequence that might be poorly processed in some heterologous expression systems. Consequently, in *Lactococcus lactis it was exch*anged for the lactococcal Exp4 signal peptide, which ensured an efficient secretion of MNaseB (24).

In this study, we compare two forms of micrococcal nuclease, MNaseA and MNaseB, and their enzymatic properties in high salt conditions. We have generated a construct design and optimized expression conditions and biochemical assaying to produce and characterize MNaseA for robust and reproducible utilization in molecular biology methods that require the presence of high salt concentrations. Furthermore, we compared purified MNaseA to a commercially available nuclease and, as a proof-of-concept, applied it to Ribo-seq of the halophilic model archaeon *H. volcanii, where it* matches and exceeds the performance of its commercial counterpart. Consequently, this work provides a simple protocol for the efficient production of highly active MNaseA, making future Ribo-seq studies of halophilic organisms more accessible.

## RESULTS

### MNase construct design and expression optimization

Previously, both the mature and the propeptide forms of microccocal nuclease (MNaseA and MNaseB, respectively) have been produced and/or utilized as tools in molecular biology (13, 22). Although the enzymatic activity of both MNase forms have been characterized in standard reaction conditions, the extent to which high salt concentrations, as encountered in applications involving e.g. halophilic archaea, impacts on MNase activity remains to be determined. To address this, we set out to produce and compare the activity of both MNase forms in high salt containing reactions. To prepare the MNase constructs (Supp. Fig. S1), we first extracted the MNaseB sequence from the *nuc gene of S. aureus (Genbank no*. V01281.1) and designed a codon optimized MNaseB synthetic gene construct with 6×His-tag on its C-terminus to allow purification using Ni-NTA affinity resin (Supp. Fig. S1A,B and Experimental procedures). Furthermore, inspired by previous MNase constructs (21, 22), we added a 21 amino acids (aa) long *E. coli OmpA signal*ing sequence to the N-terminus of MNaseB, thus targeting the produced protein to the periplasm and decreasing its toxicity to the expression host (Supp. Fig. S1B and Supp. Table S1). Based on the known OmpA signaling peptide cleavage mechanism, we expected the MNases to be expressed as OmpA containing proteins, followed by translocation to the periplasm and cleavage by signal peptidase I (25). To evaluate the expression conditions, we calculated the expected molecular weights of OmpA-containing proteins and their processed forms (Fig. 1A). Moreover, we confirmed the likelihood of OmpA signal peptide cleavage using SignalP 6.0, a protein language model-based tool (26) (Supp. Fig. S2A, Supp. Table S2). As expected, for both OmpA-fused MNases, the cleavage sites were predicted to be between 21-22 aa (probability >0.97) with secretory pathway signal peptidase I (SP(Sec/SPI)) being the main peptidase (probability >0.99) involved in signal peptide cleavage (Supp. Table S2). Consequently, we cloned the MNaseB synthetic gene construct to a pET28a backbone (Supp. Fig. S1C,D), from which the MNaseA construct was generated by deleting 19 AA from the N-terminus of MNaseB (see Experimental procedures).

**FIGURE 1.**
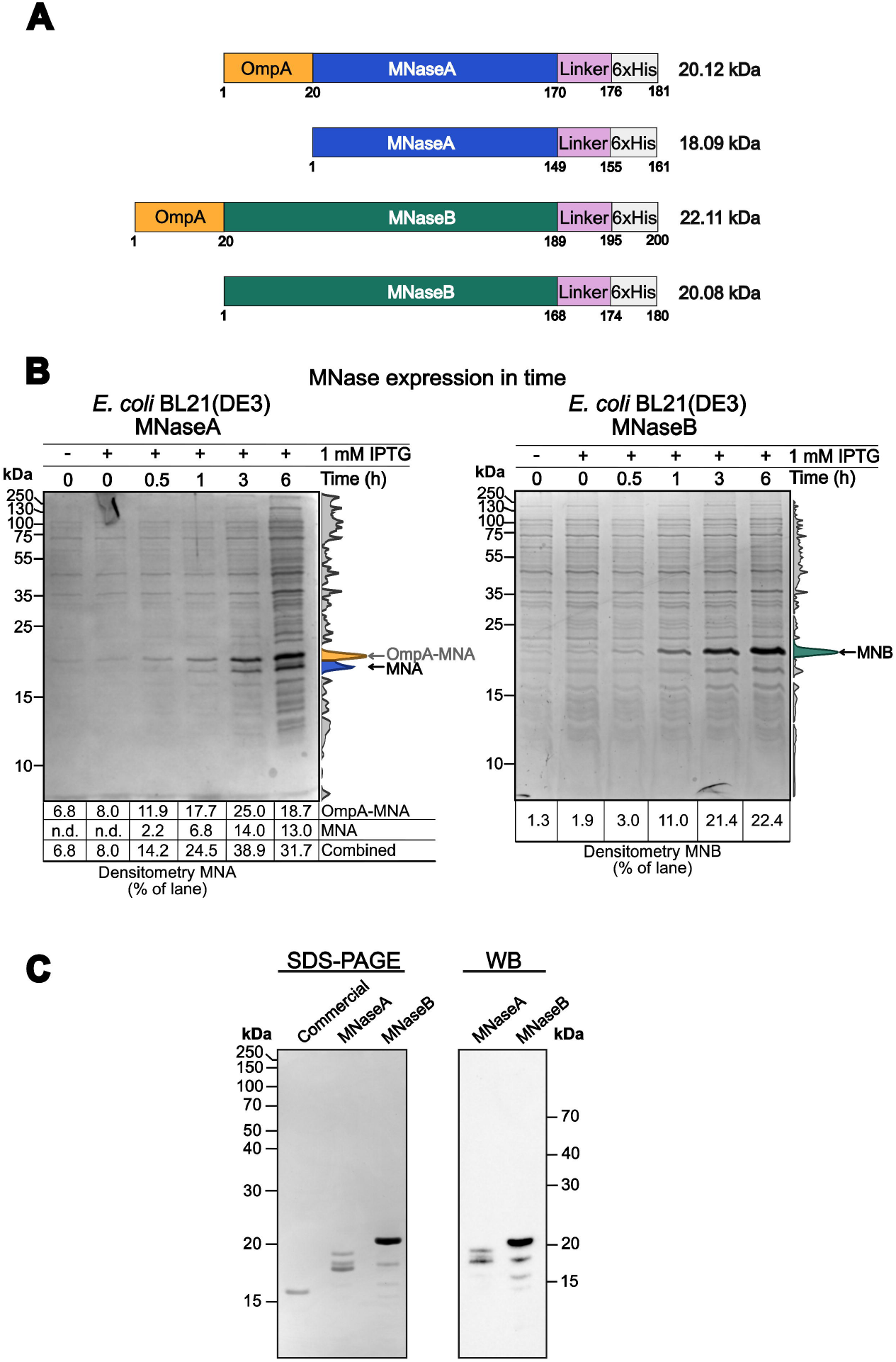
**(A)** Schematic representation of MNase forms produced from expression vectors. MNaseA (MNA) and MNaseB (MNB) are expressed with OmpA. The OmpA signaling peptide is cleaved endogenously during translocation of MNase to periplasm. **(B)** Expression of MNaseA and MNaseB at different time points with densitometrical quantification. MNaseA partially retains the OmpA peptide, while MNaseB is only expressed without the OmpA peptide. **(C)** Comparison of purified MNases with commercially available MNase on SDS-PAGE. The identity of purified MNases is confirmed by western blot with anti-6×His antibodies.

Next, we assessed protein expression from the MNaseA and MNaseB constructs as a time course experiment in the commonly available *E. coli expression* strains BL21(DE3), BL21(DE3)pLysS, and BL21(DE3)pLysE, respectively (Fig. 1B, Supp. Fig. S2B). Optimal expression for both MNases was achieved in *E. coli BL21(DE3) s*train with >20% of all protein content being MNase (Fig. 1B). MNaseA was expressed as two distinct proteins with a size of ∼18 and 20 kDa, respectively. On the other hand, MNaseB was expressed as a distinct ∼20 kDa protein (Fig. 1B). This corresponds to the predicted size of MNaseB and was further confirmed by anti-his-tag western blot analysis of the purified protein (Fig. 1C, Supp. Fig. S2C).

Given the size of the OmpA signaling peptide (∼2 kDa), we reasoned that the 20 kDa protein corresponded to OmpA-MNaseA-6×His, whereas the 18 kDa protein was MNaseA-6×His (Fig. 1A,B). To confirm our hypothesis, we analyzed purified MNaseA (average yields 11-17 mg per 1 L expression culture) (Supp. Fig. S2C) by western blot to verify their identity (Fig. 1C). The presence of unprocessed OmpA-MNaseA-6×His might be attributed to the high protein expression level, which saturates the secretion pathway machinery and leads to inefficient cleavage of the OmpA signaling peptide. To circumvent this problem, we used an alternative expression strain, *E. coli Lemo21(DE3)*, with improved control of the T7 promoter regulation (27). This allowed us to finetune MNaseA overexpression conditions by titrating the L-rhamnose promoter, which resulted in the enrichment of processed MNaseA-6×His over the unprocessed OmpA-containing form albeit with slightly decreased overall expression (∼13.4% of whole cell protein content) (Supp. Methods, Supp. Fig. S2D).

### MNaseA was more active in high salt concentration reactions than MNaseB

To ensure reproducible production and to limit batch-to-batch variation, we set up a kinetic activity assay based on absorbance measurement at 260 nm (A_260_) using optimal reaction conditions over a range of MNase concentrations (Fig. 2A, Experimental procedures). First, we benchmarked our specific activity assay with a commercially available nuclease to ensure its reproducibility (Supp. Fig. S3A). Second, using this assay, we compared our MNases produced in *E. coli BL21(DE3) t*o the commercial nuclease and calculated the specific activity for each MNase (Fig. 2B). The specific activities of the produced MNases (18.1 ± 3.6 U/µg for MNaseA and 26.3 ± 6.2 U/µg for MNaseB) were comparable to the commercial nuclease (15.2 ± 3.0 U/µg) (Fig. 2B). Additionally, we could not determine any difference in activity at optimal reaction conditions between the unprocessed (OmpA-containing) or processed form of MNaseA (Supp. Fig. S3B).

**FIGURE 2.**
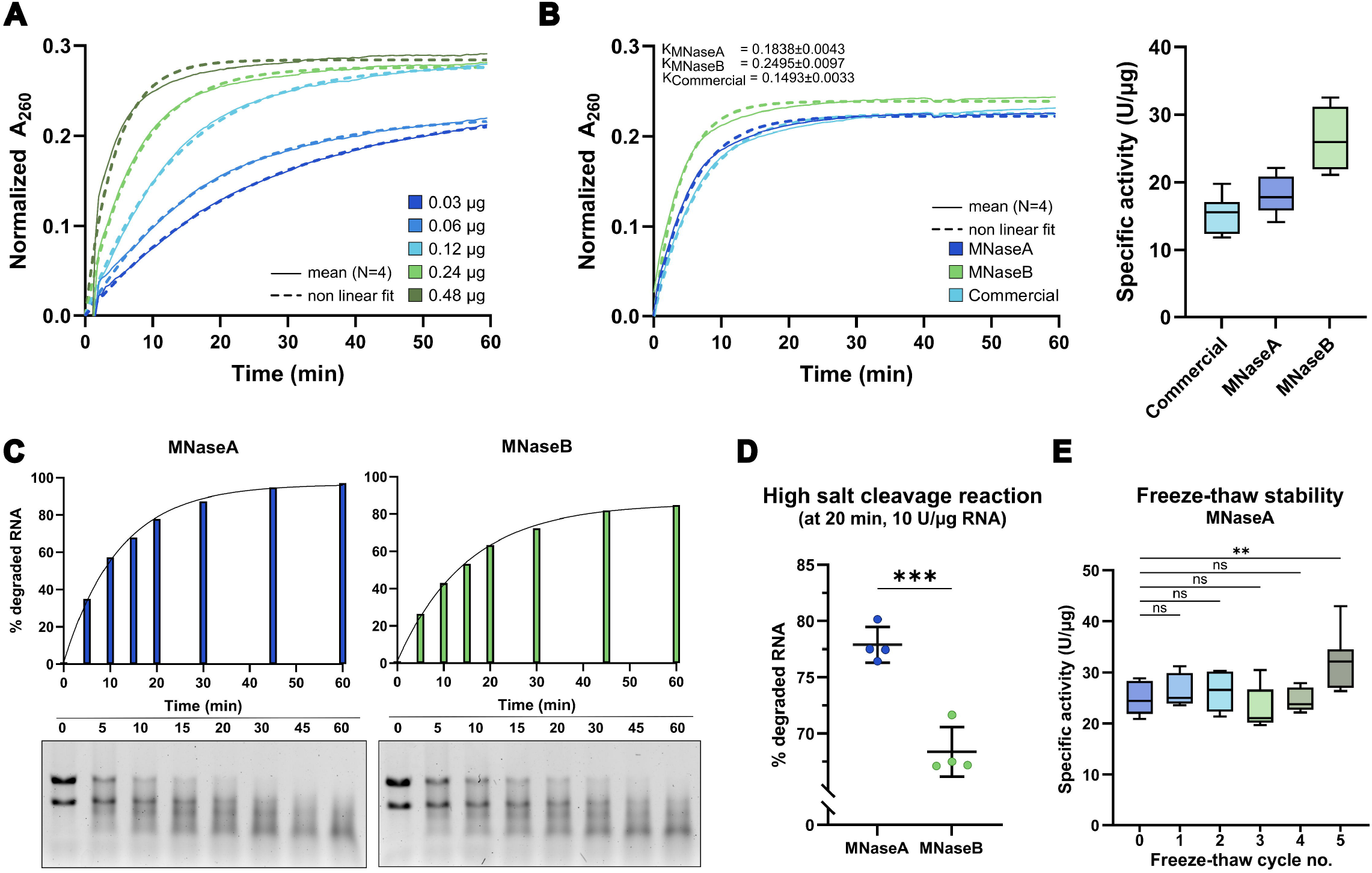
**(A)** Typical output of activity assay to measure specific activity of MNases. The assay was performed as a kinetic measurement with serial dilutions of the enzyme (mean; N=4). **(B)** Specific activities of MNases in optimal conditions. Activity was measured as a kinetic measurement (left panel; mean, N=4) and specific activity (Tukey box plot, N=8) was expressed as units per µg of protein (U/µg). K shows the reaction rate constants for each enzyme. **(C)** Performance of MNaseA and B in high salt reactions over time. MNaseA was more active in high salt than MNaseB. A representative (N=1) densitometrical analysis (top panel) of MNase activities expressed as % of degraded RNA at time 0 with gels (bottom panel) of each time point. **(D)** Densitometrical quantification of MNaseA and B activity in high salt reactions at 20 min. Samples were compared by unpaired two-tailed Student’s t-test (N=4; *** p=0.0004). (E) Stability of MNaseA stored at -80 °C during freeze-thaw cycles. The activity was measured as kinetic measurement and then specific activity of MNase was calculated after each cycle. Samples were compared using ordinary one-way ANOVA (N = 8; ns – not significant; ** p=0.0028). Error bars in all panels represent standard deviation

Next, we compared the nuclease activity of the produced MNases in high salt reaction conditions to mimic the ribosome footprinting reactions used for haloarchaea, which require >3 M KCl (18, 28). However, we could not utilize the aforementioned kinetic activity assay (A_260_ measurement) due to a known incompatibility with high salt concentrations (29). Therefore, we established a simple RNA cleavage assay in high salt reaction conditions using 10U of nuclease per µg RNA followed by gel analysis and densitometric image quantification (Fig. 2C, Experimental procedures). As expected, both MNaseA and MNaseB cleaved RNA, although MNaseA had a slightly higher activity (Fig. 2D). This was further confirmed by testing various nuclease-to-RNA ratios and different reaction time points (Supp. Fig. S3C). Taken together, MNaseA outperformed MNaseB in high salinity reactions, even though its specific activity (U/µg) was lower in optimal reaction conditions (Fig. 2B-D). We also investigated whether OmpA has an inhibitory effect on the activity of MNaseA by comparing unprocessed (OmpA-containing) and processed forms of MNaseA in high salt reactions (Supp. Fig. 3D). Similarly to assays performed in optimal reactions conditions (Supp. Fig. 3B), we could not detect any significant difference in enzymatic activity between the processed and unprocessed forms (Supp. Fig. 3D). Therefore, we decided to use MNaseA produced in *E. coli BL21(DE3) f*or further downstream applications in haloarchaeal ribosome profiling.

**FIGURE 3.**
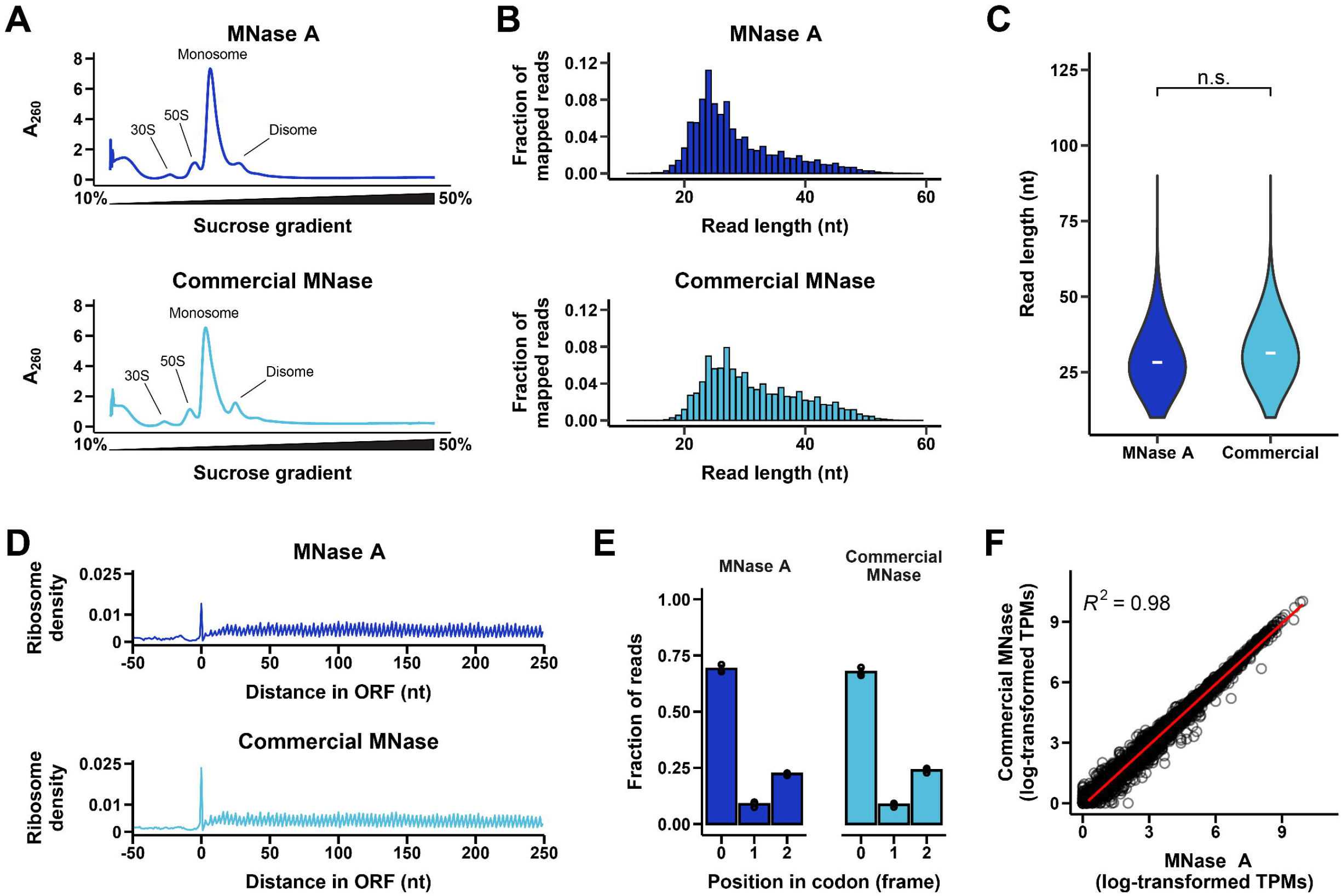
Comparison of ribosome profiling data generated using either MNase A (dark blue) or commercial MNase (light blue) to generate ribosome-protected footprints. **(A)** Absorbance traces (A_260_) of ribosome fractions along a 10-50% sucrose gradient. Treatment with MNase collapsed polysomes into monosomes. **(B)** Ribosome footprint read length distributions as bar graphs; reads per length were normalized to the total number of genome-mapped reads. **(C)** Read length distributions as violin plots. Distribution means marked by (−). Statistical comparison between distributions in each biological replicate showed no significant difference (permutation test with distribution mean as test statistic, 10,000 resamples, independent; p-value = 0.6653). **(D)** Metagene plots showing the mean ribosome density on all genes aligned at their start codon using 5’-end mapped reads. **(E)** Fraction of elongation-associated 27 nt reads mapping to each position within codons (0, +1, and +2). Bars show mean frame positions across three biological replicates; individual biological replicates are represented by points. **(F)** Correlation plot of ribosome occupancy per gene between samples treated with MNase A and commercial MNase within a biological replicate. One out of three biological replicates for each enzyme is shown in (A-D) and (F).

### MNaseA retained its activity following multiple freeze-thaw cycles

To determine if MNaseA could be used routinely in a reproducible manner, we examined its activity over the course of multiple freeze-thaw cycles. To this end, we formulated two MNaseA storage buffers – one containing 5% glycerol and another supplemented with 50% glycerol. Next, we performed five cycles of freezing at the storage temperature (−80 °C) and thawing on ice while examining the activity of the MNase during each cycle (Fig. 2E, Supp. Fig. S3D). MNaseA retains >90% of its activity after five freeze-thaw cycles when stored in 5% glycerol (Fig. 2E), whereas >80% of the activity remains when stored in 50% glycerol (Supp. Fig. S3D). We also examined how MNaseA supplemented in 50% glycerol stores at -20 °C as it is often the preferred storage temperature and prevents freeze-thaw stress of the enzyme (30). Despite the presence of 50% glycerol as a cryopreservative, this storage temperature proved to be less optimal, resulting in an almost 40% decrease in MNaseA activity (Supp. Fig S3E).

### MNaseA was suitable for ribosome profiling in haloarchaea

In ribosome profiling experiments, ribosomes and the associated mRNAs are harvested, treated with an endonuclease to digest mRNA not protected by the ribosome, and fractionated on a sucrose gradient (14, 18) (Supp. Fig. S4). The RNA (ribosome footprints) is then isolated from the monosome fractions of the sucrose gradient, used for library preparation, and deep sequenced. To assess the suitability of the purified MNaseA for ribosome footprinting in haloarchaea, we prepared Ribo-seq libraries for *H. volcanii ribosome pe*llets with MNaseA and with a commercial nuclease (Fig. 3).

We used a high-salt buffer (see Experimental procedures) as previously described for *H. volcanii (18), to mi*nimize ribosome splitting during harvesting and fractionation. This was essential because *H. volcanii has an intr*acellular salt concentration of ∼2 M KCl (18). To ensure efficient nuclease activity under those high salt conditions, we used a nuclease:RNA ratio of 250 U nuclease/1 µg RNA (RNA concentrations measured by Qubit) for both MNaseA and the commercial nuclease.

Following digestion of ribosome pellets and fractionation, we observed high A_260_ absorbance of monosome-associated sucrose gradient fractions and low absorbance of polysome-associated fractions with both nucleases (Fig. 3A). Small disome (2 monosomes) peaks were also present after treatment; these may correspond to naturally occurring endogenous ribosome collisions, which are known to be resistant to nuclease digestion (31). After library construction, quality control, and sequencing (Supp. Fig. S5), we analyzed the mapped ribosome-protected footprint length distributions for each nuclease (Fig. 3B, Supp. Fig. S6). MNaseA produced ribosome footprint lengths comparable to that of the commercial nuclease (Fig. 3B). There was no statistical difference between read length distributions of digestions performed with MNaseA or the commercial nuclease across all biological replicates (Fig. 3C, Supp. Fig. S6), indicating that digestion by MNaseA and the commercial nuclease produced comparable ribosome footprints.

Hallmarks of elongating ribosomes are high ribosome occupancy in ORFs and a 3-nucleotide periodicity in ribosome footprints, corresponding to the ribosome moving one codon at a time along the mRNA (14, 17, 18).

Using metagene plots, which average ribosome density across numerous ORFs aligned at their start codons, we found high ribosome density in ORFs and relatively low density in the untranslated region upstream of the start codon for both enzymes (Fig. 3D). We also found a similar 3 nt periodicity in translated regions (Fig. 3D). These results were consistent regardless of whether ribosome density was calculated based on the 5’-end (Fig. 3D) or 3’-end (Supp. Fig. S7) of the ribosome footprint. At the sub-codon scale, we showed that both enzymes produced ribosome footprints that mapped predominantly to the first codon position at their 5’ end (Fig. 3E). This is further evidence of 3 nt periodicity and consistent with our previous observations in *H. volcanii (18). In co*nclusion, both MNaseA and the commercial nuclease produced similar ribosome profiling data that displayed signs of actively elongating ribosomes.

Finally, we compared ribosome occupancy of individual genes between libraries prepared from either MNaseA or the commercial nuclease by converting ribosome profiling reads to transcripts per million (TPMs). Translation levels of genes were highly correlated between MNaseA and the commercial nuclease (R^2^ > 0.97 for all replicates; Fig. 3F, Supp. Fig. S8), indicating that both nucleases produced comparable information about the active translatome in *H. volcanii*.

## DISCUSSION

*In this stu*dy, we described the design and purification of the *S. aureus nuclease, M*Nase, for use in high salt reaction conditions, such as Ribo-seq in halophilic organisms. Based on our experiments and in agreement with published literature, MNaseB, the precursor of MNaseA, is easier to produce thanks to the 19 aa propeptide on its N-terminus, which is known to enhance processing and secretion of the nuclease to the periplasm (20). On the other hand, MNaseA is produced as two protein products of 18 kDa and 20 kDa, respectively. The smaller protein corresponds to the fully processed form of MNaseA, where the OmpA signaling peptide has been cleaved off upon translocation to the periplasm, whereas the bigger protein represents the unprocessed OmpA-MNaseA (Fig. 1B, C). The incomplete processing and excretion of MNaseA has also been observed in previous studies (20, 21). Although the 19 aa propeptide from MNaseB is not necessary for correct folding and activity of the nuclease (21, 23, 32), previous studies suggest it is required for anchoring the protein to the cell wall (33). Additionally, Suciu and Inouye suggested that MNaseB membrane insertion is SecA independent and that during initial membrane insertion, the 19 aa propeptide counteracts the inhibitory effect of the positively charged N-terminus of MNaseA (20). Hence, the higher excretion of MNaseB can be attributed to the biochemical properties of the 19 aa propeptide, which was absent in MNaseA. To date, several strategies to enhance OmpA signal peptide cleavage and MNaseA excretion have been applied. For example, it has been suggested that excretion enhancing alterations, such as those conferred by the MNaseA double mutant F15W-W140F may be applied (20). However, this double mutation also affects the structural stability of the enzyme (20, 34) and its effect on MNase activity has not been extensively studied. Other approaches employ expression and excretion of MNaseA in alternative expression hosts using excretion vectors, such as *Corynebacterium glutamicum, Bacillus subtillis or Lactobacillus lactis, where MNa*seA can be purified directly from media (23, 24, 32). Such excretion systems are scalable and amendable for bioreactor production. Despite these advantages, the resulting nuclease activity remains too low—2000 U/mg protein in *L. lactis (24)—* to be applicable for high salt conditions, such as in methods for halophilic organisms, where small volume and over 250 U of MNase/μg of RNA is necessary in Ribo-seq experiments (18). Moreover, such expression hosts are not commonly available in research laboratories, which might further diminish their appeal. To circumvent these problems, we developed an optimized production protocol for MNaseA in the *E. coli Lemo21(DE3)* strain (27). Thanks to the improved control of the T7 promoter through rhamnose titration, we obtained a significantly higher portion of processed MNaseA, which offset the slight decrease in total MNaseA yield caused by an overall reduction in expression (Supp. Fig. S2D).

To our knowledge, MNaseA and MNaseB have not previously been directly compared in terms of their biochemical properties and/or enzymatic activities. Therefore, we compared these two MNase forms using the same amount of activity units on the same substrate. While MNaseA and MNaseB performed comparably in optimal reaction conditions (low salt concentration and pH 8), MNaseA exhibited a higher nuclease activity in high salt conditions (>3M KCl and pH 7.5). As both forms of the nuclease were otherwise identical, it stands to reason that the observed differences were likely to arise from the biochemical properties of the 19 aa propeptide sequence of MNaseB, which similarly to the OmpA signaling peptide has a mostly neutral charge (20). Consequently, purified MNaseA may also be partially affected by this phenomenon, as it contains a mixture of both processed and unprocessed (i.e. OmpA-containing) nuclease. However, the amount of unprocessed OmpA-MNaseA in final purified enzyme was minor (Fig. 1C) and therefore, was not likely to have a significant impact. Moreover, MnaseA production in *E. coli Lemo21(DE3)* resulted in fully processed MNaseA, which allowed us to evaluate and confirm that comparable enzymatic activities were obtained regardless of the presence of OmpA (Supp. Fig. 2B, D). When characterizing the properties of MNaseA, we observed that it preserved well at -80 °C when stored in a buffer supplemented with merely 5% glycerol rather than with 50% glycerol. Generally, protein cryopreservation is achieved by the addition of cryoprotectants (e.g. 25-50% glycerol), which inhibit freeze/thaw induced stress, such as protein aggregation and ice crystal formation, thereby protecting the native fold (35, 36). However, a recent study shows that hydroxyperoxide lyase retained more activity when stored in 10% glycerol at -80 °C, whereas its residual activity decreased rapidly during freeze/thaw cycling when the enzyme was stored in 40% glycerol, suggesting the need for empirical testing of the optimal glycerol concentration (37). Alternatively, the reduced stability of MNaseA in the presence of 50% glycerol might be attributed to a dilution effect, as an increase in glycerol lowers the protein concentration (38, 39). Correspondingly, our stability assaying of MNaseA stored at -20 °C showed a similar decrease in activity, suggesting that freeze/thaw stability might be improved by the addition of other enzyme stabilizing additives (40). Nevertheless, we concluded that the optimal storage condition for MNaseA is at -80 °C with 5% glycerol, although a suitable storage buffer for -20 °C might be obtained through further optimization.

Previously, purified MNaseB was successfully used for Ribo-seq in mesophilic bacteria, namely *E. coli, showing s*imilar properties to the commercial counterpart (22). Nonetheless, neither MNaseA nor MNaseB have been previously utilized for similar methods in halophilic organisms. Since MNaseB showed a lower activity in high salt reactions, we decided to proceed only with MNaseA and compare its ribosome footprinting capabilities to those of a commercial nuclease by performing Ribo-seq on the halophilic model organism *H. volcanii. Indeed, M*NaseA produced ribosome footprints of similar length and showed an expected periodicity that reflects the ribosome movement on mRNA (Fig. 3). Taken together, our data shows that MNaseA produced with our construct design and expression conditions is on par with or even slightly exceeds the activity obtained by the commercial nuclease, especially in high salt reactions. Importantly, the protocol is robust and easily reproducible, allowing investigators with access to basic laboratory facilities to economically (at ∼25% cost of commercial enzyme/µg of enzyme) produce a highly active nuclease.

An economic source of nuclease is especially critical for ribosome profiling in halophilic organisms. To prevent ribosome splitting during ribosome harvest and fractionation, a high-salt buffer is necessary, which in turn requires high amounts of nuclease for proper RNA digestion. In our experiments, we used 250 U nuclease/1 µg RNA, with RNA concentrations measured fluorometrically. As previous ribosome profiling experiments rely on absorbance measurement to quantify RNA, a correction factor is required for comparison to our protocol (see Table S3 and S4). Thus, the nuclease:RNA ratio used here (22 U nuclease/1 µg RNA; approximated as a absorbance-based concentration) was approximately two-fold higher than the ratio previously reported for ribosome profiling in *E. coli (∼10* U nuclease/1 µg RNA) (17), where salinity is not an issue. However, when compared to previous experiments with *H. volcanii (∼50* U nuclease/1 µg RNA) (18), we used slightly less MNase while obtaining an efficient digestion (Figure 3). Importantly, it is worth noting that further reducing the nuclease:RNA ratio might result in residual polysome peaks, indicative of incomplete digestion of unprotected RNA, whereas lowering the amount of RNA increases the risk of library preparation failure. Consequently, the amount of nuclease required for ribosome footprinting in *H. volcanii and similar* organisms remains relatively high, but the cost of this enzyme has been lessened by the work presented here.

## EXPERIMENTAL PROCEDURES

### Preparation of MNase constructs

The complete MNase coding sequence was extracted from the *nuc gene (966 n*t) of *S. aureus (GeneBank n*o. V01281.1) (Supp. Fig. S1). The *nucB precursor (*MNaseB) is encoded by residues 394-909 (19.1 kDa), whereas *nucA (MNaseA), t*he fully processed form, is encoded by residues 463-909 (16.8 kDa).

The MNaseB construct was ordered as synthetic gene (Genewiz) containing a T7 promoter, lac operon, ribosome binding site (RBS), and OmpA signaling sequence (MKKTAIAIAVALAGFATVAQA) at the N-terminus of the *E. coli codon optim*ized *nucB gene, follo*wed by a 5 aa linker. Codon optimization was done with IDT’s Codon Optimization Tool (https://www.idtdna.com/CodonOpt). This synthetic gene was subsequently re-cloned into the expression vector pET28a(+) (Novagen), containing an in-frame 6×His tag at the C-terminus, between the *BglII and XhoI* sites. The final construct was propagated in *E. coli DH5α* in LB-Miller (LB) growth medium (10 g/L tryptone, 5 g/L yeast extract, 10 g/L NaCl) supplemented with 50 µg/mL kanamycin. For a detailed vector map of pET28a(+)-T7-OmpA-MnaseB-6×His and the complete sequence of the synthetic gene, see Supp. Fig. S1 and Supp. Table S1, respectively.

The MNaseA expression vector (pET28a(+)-T7-OmpA-MnaseA-6×His) was generated by deleting the first 57 nt (19 aa) from the *nucB gene in the* pET28a(+)-T7-OmpA-MnaseB-6×His construct using site-directed mutagenesis (41) with non-overlapping mutagenic primers (Supp. Table S1). For mutagenesis protocol, please refer to Supporting Methods.

The sequences of both expression vectors were verified by Sanger sequencing (Eurofins) and are available from Addgene (#214808 and #214809).

### Expression and purification of MNaseA

MNaseA was expressed in *E. coli BL21(DE3) f*rom pET28a(+)-OmpA-MnaseA. Freshly transformed *E. coli BL21(DE3) c*ells were inoculated to LB supplemented with 50 µg/mL kanamycin and grown overnight at 37 °C with shaking at 200 rpm. The stationary cultures were diluted to OD_600_ = 0.1 and grown to an OD_600_ of 0.5-0.6, at which point protein expression was induced by IPTG (ThermoFisher Scientific, #R0392) addition (1 mM final concentration) and subsequently carried out at 30 °C with shaking at 200 rpm for 6 h.

The cells were collected by centrifugation at 5,000 x *g for 15 min* at 4 °C, and the pellets were weighed and resuspended in 2.5-3 mL lysis buffer per 1 g of wet cell pellet. The composition of lysis buffers are as follows: MNaseA lysis buffer LB-MNA [50 mM Tris-HCl (pH 7.5), 250 mM NaCl, 1 mM CaCl_2_, 5 mM imidazole (pH 7.5), 5% (vol/vol) glycerol, 1× Halt protease inhibitor (ThermoFisher Scientific #87786)]. To enhance subsequent lysis by cryomilling, the resuspended culture was dispensed into liquid nitrogen using a serological pipette. The frozen pellet droplets were transferred to 50 mL conical tubes and stored at -80 °C.

Frozen pellet droplets were lysed in 6870 Large Freezer/Mill (Spex) at 8 Hz for 5 cycles (1 min grinding, 1 min cooling). Lysates were transferred to 50 mL conical tubes and allowed to melt on ice. Once melted, an additional 2.5-3 mL of the respective lysis buffer per 1 g wet cell pellet was added to the lysate. The lysate was mixed by inverting and incubated on ice for 10-20 min. Cell debris were removed by centrifugation at 10,000 x *g, 30 min, +*4 °C. The clarified lysate (supernatant) was collected and mixed with 2-3 mL 50% slurry of Ni-NTA resin beads (ThermoFisher Scientific, #88221) washed twice in LB-MNA. The lysate was then incubated with Ni-NTA beads for 1-2 h at 4 °C, slowly rotating on tube revolver rotator. The lysate mix was loaded onto a gravity flow column (ThermoFisher Scientific, #89896) and washed with 2 column volumes (CV) of lysis buffer, followed by 10 CV of wash buffer [50 mM Tris-HCl (pH 7.5), 250 mM NaCl, 20 mM imidazole (pH 7.5) and 5% (vol/vol) glycerol]. The MNase was eluted from Ni-NTA beads by 6 CV of elution buffer [50 mM Tris-HCl (pH 7.5), 250 mM NaCl, 250 mM imidazole (pH 7.5) and 5% (vol/vol) glycerol]. All fractions were stored at 4 °C until analyzed by SDS-PAGE. The protein concentrations were measured by Bradford assay (42).

Fractions were analyzed on 5-15% discontinuous (5% stacking/15% resolving) SDS-PAGE gels and the eluted fractions containing MNase were pooled together and concentrated on a 10K MWCO protein spin concentrator (ThermoFisher Scientific; #88513). Buffer exchange was done by diluting the concentrated protein with Storage Buffer [50 mM Tris (pH 7.5), 50 mM NaCl, 1 mM EDTA (pH 8.0), and 5% (vol/vol) glycerol] and concentrating the protein again. This was repeated 5 times to eliminate any residual imidazole from the final MNase preparation. MNase was then aliquoted and stored at -80 °C. The purity of MNase was confirmed on 5-15% discontinuous SDS-PAGE gel. To store enzyme at -20 °C, the final protein was supplemented with 50% glycerol.

For buffer composition for MNaseB purification, description of alternative lysis methods (sonication) and MNaseA expression in *E. coli Lemo21(DE3)*, see Supporting Methods.

### Western blot analyses

100 ng of each protein was separated on 5-15% discontinuous SDS-PAGE gel. After electrophoresis, gel was equilibrated in Bjerrum Schafer-Nielsen transfer buffer (48 mM tris, 39 mM glycine, 20% methanol) and proteins were transferred to PVDF membrane using Trans-Blot SD transfer cell (Biorad). After transfer, membrane was blocked in 5% milk in PBST (1xPBS, 0.05% Tween-20) for 1 h followed by incubation with 1:2,000 diluted 6x-His tag monoclonal antibody (produced in mouse; Invitrogen; #MA1-21315) at 4 °C, overnight. Next day, the membrane was washed 3× in PBST and incubated 30 min at room temperature with anti-mouse HRP-antibody, diluted 1:10,000 (Cytiva; #NA931) in 5% milk in PBST. After incubation, the membrane was again washed 3× in PBST and the proteins were detected by chemiluminescence using ECL Western Blotting Substrate (ThermoFisher Scientific; #32209) according to manufacturer’s instructions. The chemiluminescence was recorded by ChemiDoc (Biorad).

### Activity assays

#### Kinetic measurement of specific activity and unit definition

The MNase specific activity was measured as previously described with a few modifications (22). The reactions were prepared by mixing 50 µL substrate (0.2 mg/mL dsDNA in 10 mM Tris-HCl pH 8.0 and 10 mM CaCl_2_) on a 96-well UV-compatible plate with 50 µL MNase dilution in 10 mM Tris-HCl pH 8.0. In each measurement, serial dilutions of MNase were assayed (0.03, 0.06, 0.12, 0.24, and 0.48 µg). The reactions were immediately placed into a pre-heated plate reader (Enspire, PerkinElmer) and reactions were monitored for 60 min at 25 °C, with repeat A_260_ absorbance measurement every 30 s. Data points were fitted to an exponential equation with one-phase association using GraphPad Prism (v10.1.0) and the following formula *A*_*x*_ = *A*_*0*_ + (*P-A*_*0*_*) × (1 – e*^(−*K* × *x*)^) (*A*_*x*_ : *absorptio*n A_260_ at time x; *A0: absorptio*n A_260_ at time 0; *P: plateau =* absorption A_260_ at infinite time; *K: rate cons*tant as reciprocal time; x: time). The specific activity was calculated from initial slopes for time 0 and 1 min. One unit (U) is defined as an increase of 0.005 A_260_ units per 1 min.

#### Comparison of activity in reactions with high salt concentration

The enzymatic activity of purified MNases in high salinity was compared by cleavage of total RNA and subsequent analysis on 1% denaturing agarose gels. Reactions were prepared by diluting total RNA in 1× lysis buffer (3.4 M KCl, 100 mM MgCl_2_, 50 mM CaCl_2_, 10 mM Tris-HCl pH 7.5) to concentration 250 ng/µL (28). The reactions were initiated by addition of 5 or 10 U MNase/µg of RNA and incubated at 25 °C. 5 µL samples containing 1.25 µg RNA were collected at 0, 5, 10, 15, 20, 30, 45 or 60 min and MNase activity was quenched by addition of 20 µL formamide LD (90% formamide, 1xTBE, 0.075% Orange G) mixed with 5 µL 0.125 M EDTA. Samples were denaturated at 65 °C for 5 min and analyzed on 1% agarose gel prepared in 1xTBE. The degree of RNA degradation was quantified densitometrically using volume tool in Image Lab, ver. 6.0.0 (Biorad). The RNA densities were first normalized to the lane (background) and then the % of degraded RNA was calculated as normalized ratio of cleaved (at time x)/intact RNA (at time 0 min).

### Haloferax volcanii culture conditions

*H. volcanii DS2 strain* H26 (Δ*pyrE2) (Hv) si*ngle colonies were picked and grown overnight at 42 °C with shaking at 220 rpm (Eppendorf Innova S44i) in Hv-YPC medium (https://haloarchaea.com/wp-content/uploads/2018/10/Halohandbook_2009_v7.3mds.pdf) until saturation (OD_600_ > 1.0). The cultures were then diluted to OD_600_ = 0.01 in fresh media and grown to OD_600_ = 0.6.

### Cell lysis, ribosome pelleting, MNase digestion, and sucrose gradients for ribosome fractionation

As previously described (28), cells were harvested by dispensing culture into liquid nitrogen, forming small pellets. Batches of 50 g of pellets were supplemented with 100 μg/mL anisomycin (Sigma-Aldrich, #A9789) and pulverized in a cryomill (6870 Large Freezer/Mill (Spex)) for 8 cycles (1 min grinding at 10 Hz, 1 min cooling). Cell lysates were thawed at RT and pre-cleared at 10,000 × *g for 15 min* at 4 °C. Ribosomes were pelleted from pre-cleared lysate over a 1 M sucrose cushion [sucrose dissolved in 1× lysis buffer (3.4 M KCl, 100 mM MgCl_2_, 50 mM CaCl_2_, 10 mM Tris-HCl pH 7.5)] in an ultracentrifuge with a Type 45 Ti rotor (Beckman Coulter) at 40,000 rpm (185,511 × *g) for 3 h a*t 4 °C. Ribosome pellets were resuspended in 300 μL 1× lysis buffer, then pooled for each biological replicate. RNA concentrations of pooled pellets were measured using the Qubit RNA Broad Range assay (ThermoFisher Scientific, #Q10210), then pellets were split back into equal volumes. For each biological replicate, equal amounts of RNA were digested either with Roche MNase (Nuclease S7, Roche, #10107921001) or with MNaseA, for 1 h at 25 °C, at a ratio of 250 U per 1 μg RNA. Digested pellets were loaded onto a 10-50% sucrose gradient (sucrose dissolved in 1× lysis buffer) and ultracentrifuged using a SW41 Ti rotor (Beckman Coulter) at 40,000 rpm (273,620 × *g) for 2*.*5 h* at 4 °C. Gradients were fractionated into 450 μL fractions to resolve ribosome 30S and 50S subunits, monosomes, and polysomes. Fractions were flash-frozen on dry ice for subsequent RNA extraction.

### RNA extraction and ribosome footprint library preparation

TRIzol LS (Invitrogen) was used according to manufacturer instructions to extract RNA from sucrose gradient fractions corresponding to 70S monosomes. Library preparation was performed as previously described (28). Briefly, 1-5 µg of RNA fragments extracted from 70S monosome fractions were used to purify 10–45 nt RNA fragments by polyacrylamide gel electrophoresis on a 15% TBE-Urea gel; RNA fragments were treated with T4 polynucleotide kinase (NEB, #M0201S), ligated to the linker (NEB Universal miRNA Cloning Linker, #S13115S) using T4 RNA ligase (NEB, #M0242), purified by an Oligo Clean and Concentrator kit (Zymo Research, #D4061), and reverse transcribed with SuperScript III (Invitrogen, #18080044) using custom primers previously described (28). DNA fragments were gel purified on a 10% TBE-Urea denaturing polyacrylamide gel, circularized using CircLigase (Epicentre, CL4115K), and PCR amplified (8–12 cycles) with Phusion polymerase (NEB) using custom primers (28). PCR products were gel extracted from a native 8% TBE gel and analyzed for size and concentration using a BioAnalyzer high-sensitivity DNA kit (standard protocol) before sequencing (Novagene, Novaseq 6000, PE150). Primer sequences can be found in Supporting Table S1.

### Analysis of ribosome profiling data

All ribosome profiling data were analyzed using previously established methods and the plastid python library (17, 18, 43). In brief, reads were trimmed using trim galore v. 0.6.10, reads corresponding to rRNA and tRNA were discarded, and the remaining reads were mapped against the *Hv NCBI RefSeq* genome (taxonomy identification 2246, GCF 000025685.1, ASM2568v1), allowing two mismatches using Bowtie2 v. 2.5.3 (44). Statistics of genome-aligned reads were produced with samtools v. 1.19 (45). Distributions from commercial MNase and MNaseA were statistically compared within biological replicates by permutation test with scipy v. 1.11.3 (distribution mean as test statistic, two-sided, independent, 10,000 resamples) (46). Genome-aligned reads were then subset to include only reads that mapped to coding sequences (CDSs); these data were converted to transcripts per million (TPMs) using Stringtie v. 2.2.3 (47), log-transformed, and compared between MNaseA and the commercial nuclease for each biological replicate.

The plastid module *metagene (subprogram count) was used* to construct metagene plots. Genome-mapped reads were subset to include only reads that mapped to CDSs, then these reads were aligned at the start codon of each CDS. Ribosome density was calculated based on the mean number of either 5’- or 3’ read ends mapped at each nucleotide position 250 nt into the CDS (default calculations are performed with median values rather than means). An additional 50 nt upstream of the start codon was also included in the analysis to compare ribosome density outside of CDSs and within CDSs.

The plastid module *psite was used to* determine the *H. volcanii ribosome P-*site offset, or the distance in nucleotides between the P-site (internal to the ribosome footprint) and the 5’ end of the ribosome footprint. This analysis was performed in a manner like metagene analysis, but stratified ribosome density by read length; a consistent offset between a start or stop codon and a spike in high ribosome density (due to the rate-limiting nature of translation initiation and termination) should be evident across various read lengths. For the current work, the P-site offset was measured as the distance between the stop codon and a spike of high ribosome density shortly upstream of the stop codon; the start codon was forgone since *H. volcanii has a high* proportion of leaderless transcripts, resulting in high ribosome density precisely at the start codon and no measurable P-site offset for this feature. The P-site offset was found to be 14-17 nt for highly abundant ribosome footprint lengths associated with elongation (14 nt offset for 24 nt footprints; 17 nt offset for 27 nt footprints). A 17 nt P-site offset for 27 nt footprints was used with the plastid module *phase_by_size to calculat*e the positions of ribosome footprint 5’-ends within codons.

## DATA AVAILABILITY

Generated sequencing data are deposited on NCBI GEO under accession number GSE269836.

## ACKNOWLEDGEMENTS

The authors thank Riitta Tarkiainen and Minna Poranen for providing the pET28a(+) vector and *E. coli BL21(DE3)pL*ysS; and Eric Johnson for kind gift of *E. coli BL21(DE3)pL*ysE strain. We thank the HiLIFE Biocomplex Unit, University of Helsinki—a member of Instruct-ERIC Centre Finland, FINStruct, and Biocenter Finland—for providing access to high-speed centrifugation services, and Molecular Ecology and Systematics Laboratory, University of Helsinki for access to plate reader.

## FUNDING AND ADDITIONAL INFORMATION

This work was supported by the Novo Nordisk Foundation (grant no. NNF19OC0054454 to L.P.S.), the Research Council of Finland (grant no. 354906 to L.P.S.), the Sigrid Jusélius Foundation (grant no. 230182 to L.P.S.), and the US National Science Foundation (grant no. MCB-2034271 to J.D.R.). The Fulbright Finland Foundation is acknowledged for mobility support (to P.G.). P.G. is a fellow of the Doctoral Programme in Integrative Life Sciences. The Margolies family is acknowledged for mobility support through an educational travel grant (to M.I.).

