## Supporting Information for "Purification of micrococcal nuclease (MNase) for use in ribosomal profiling of high-salinity extremophiles"

**Running title:** Optimized purification of MNase for use in ribosomal profiling of *Haloarchaea*;

**KEYWORDS**

Micrococcal nuclease, S7, MNase, ribosome profiling, Haloarchaea, Halophiles, Archaea

**Supporting Materials and Methods**

**Site-directed mutagenesis of MNaseB to prepare MNaseA construct**

Briefly, the PCR reaction contained 0.5 µM of each primer, 0.4 mM dNTPs, 1x HF buffer, 0.02 U/µL Phusion polymerase (ThermoFisher Scientific, # F-530L) and 2 ng template. The mutagenesis was carried out using touchdown PCR: initial denaturation at 98 °C for 5 min, 20 cycles of 98 °C for 20 s, 60-50 °C (step -0.5°C/cycle) for 20 s, 72 °C for 1 min 30 s; 10 cycles of 98 °C for 20 s, 55 °C for 20 s, 72 °C for 1 min 30 s; final extension at 72 °C for 10 min and stored at 8 °C. Following the PCR, 2 µL of the unpurified PCR product was circularized by performing kinase/ligase mixture [0.5x T4 DNA ligase buffer, 0.5x T4 PNK buffer, 0.5 mM ATP, 0.5 U/µL T4 DNA Ligase (ThermoFisher Scientific, #EL0014), 1 U/µL DpnI (ThermoFisher Scientific, #FD1703), 0.5 U/µL T4 PNK (ThermoFisher Scientific, #EK0031)]. Reaction was incubated at 22 °C for 2 h, followed by inactivation at 70 °C for 5 min and transformed to chemically competent *E. coli* DH5α. The colonies were screened by colony PCR and propagated in *E. coli* DH5α in LB supplemented with 50 µg/mL kanamycin. Sequence was verified by Sanger sequencing (Eurofins).

**Cell lysis using sonication**

As an alternative to cell lysis by cryomill, the sonication was equally efficient lysis method. For sonication, harvested cells were snap frozen as wet pellets, weighted, and stored at -80 °C until further processed. Then, the pellets were melted on ice and 5 mL of Lysis buffer was added per 1 g of wet pellet and cells were resuspended by pipetting. Sonication (Sonicator UP400 S Dr. Hielscher) was done using the following program: 10 cycles, 60% amplitude, 0.5 cycle, 20 s ON/1 min OFF. Lysate was centrifuged for 10,000 x *g,* 30 min, 4 °C. The resulting supernatant was used for purification as described for MNaseA.

**MNase B purification**

The purification is performed in same manner as for MNase A. However, the purification buffers have slightly different composition. The composition of these buffers is below:

- MNase B lysis buffer (**LB-MNB**) [50 mM Tris-HCl (pH 7.5), 500 mM NaCl, 1 mM $CaCl_2$, 5 mM imidazole (pH 7.5), 5% (vol/vol) glycerol, 1× Halt protease inhibitor (ThermoFisher Scientific #87786)]
- MNase B wash buffer (**WB-MNB**) [50 mM Tris-HCl (pH 7.5), 500 mM NaCl, 10 mM imidazole (pH 7.5) and 5% (vol/vol) glycerol]
- MNase B elution buffer (**EB-MNB**) [50 mM Tris-HCl (pH 7.5), 500 mM NaCl, 250 mM imidazole (pH 7.5) and 5% (vol/vol) glycerol]

**MNase A expression in *E. coli* Lemo21(DE3)**

To increase the OmpA, MNaseA can be also expressed in *E. coli* Lemo21(DE3) using pET28a(+)-OmpA-MnaseA plasmid. For expression, freshly transformed *E. coli* Lemo21(DE3) cells were inoculated to LB-Luria (10 g/L tryptone, 5 g/L yeast extract, 0.5 g/L NaCl supplemented with 50 µg/mL kanamycin, 35 µg/mL chloramphenicol) and grown overnight at 37 °C with shaking at 200 rpm. The stationary cultures were diluted to $OD_{600}$ = 0.1, supplemented with antibiotics and 500 µM L-rhamnose and grown to an $OD_{600}$ of 0.4-0.5, at which point protein expression was induced by IPTG (ThermoFisher Scientific, #R0392) addition (1 mM final concentration) and subsequently carried out at 30 °C with shaking at 200 rpm for 9-10 h. The cells were collected by centrifugation at 5,000 x *g* for 15 min at 4 °C and processed as described in Methods section.

**Supporting Figures**

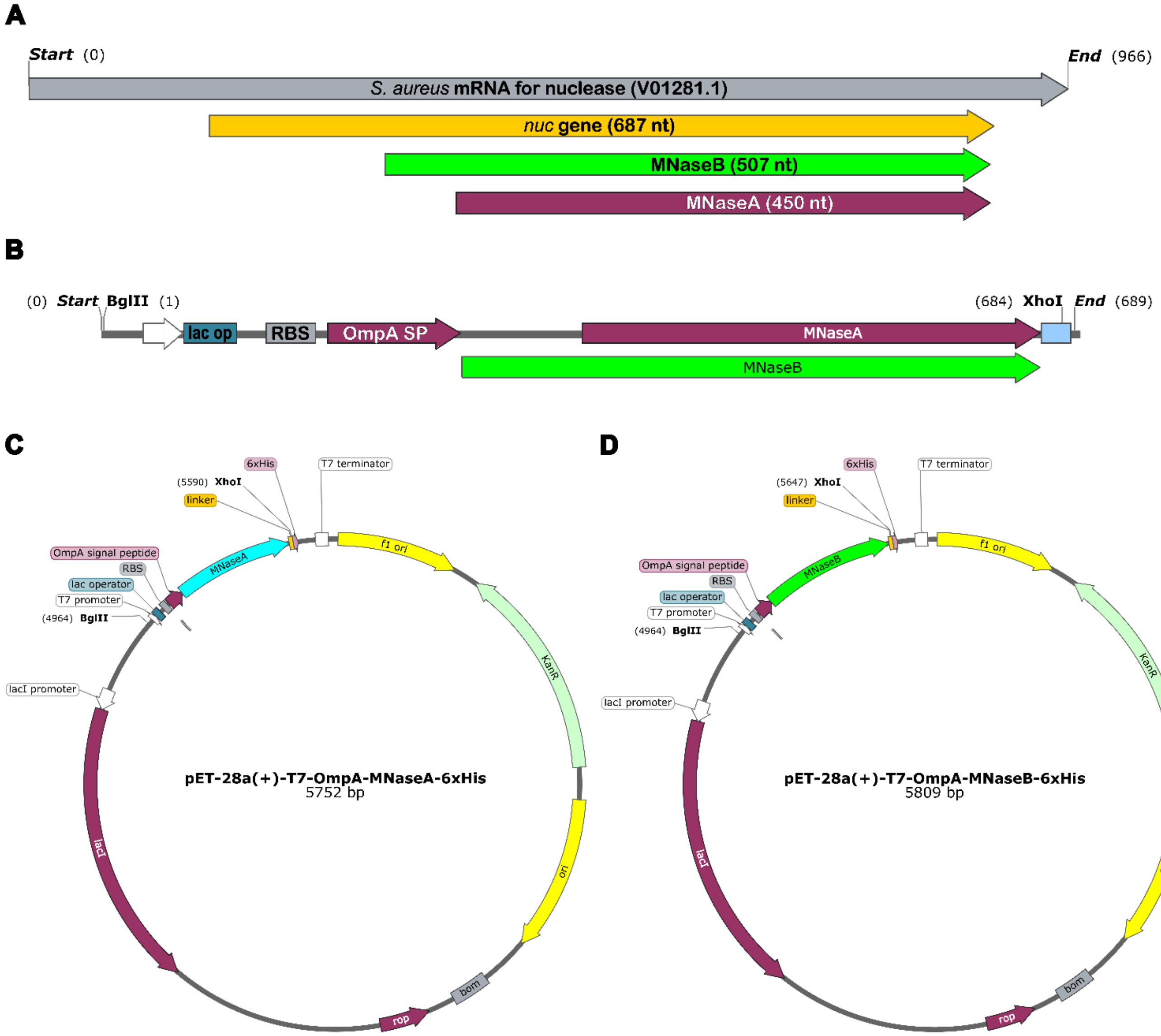


**Supporting Figure S1. (A)** Schematic view of *S. aureus* mRNA for nuclease (966 nt) organization (GenBank no. V01281.1). *Nuc* gene is encoded by 687 nt (226-912 nt) of which MNaseB is 507 nt (403-912 nt) and MNaseA is 450 nt (463-912 nt). **(B)** Schematic view of synthetic gene construct for MNase production. Plasmids designed in the study. **(C)** Plasmid for expression of MNaseA and **(D)** MNaseB, respectively.

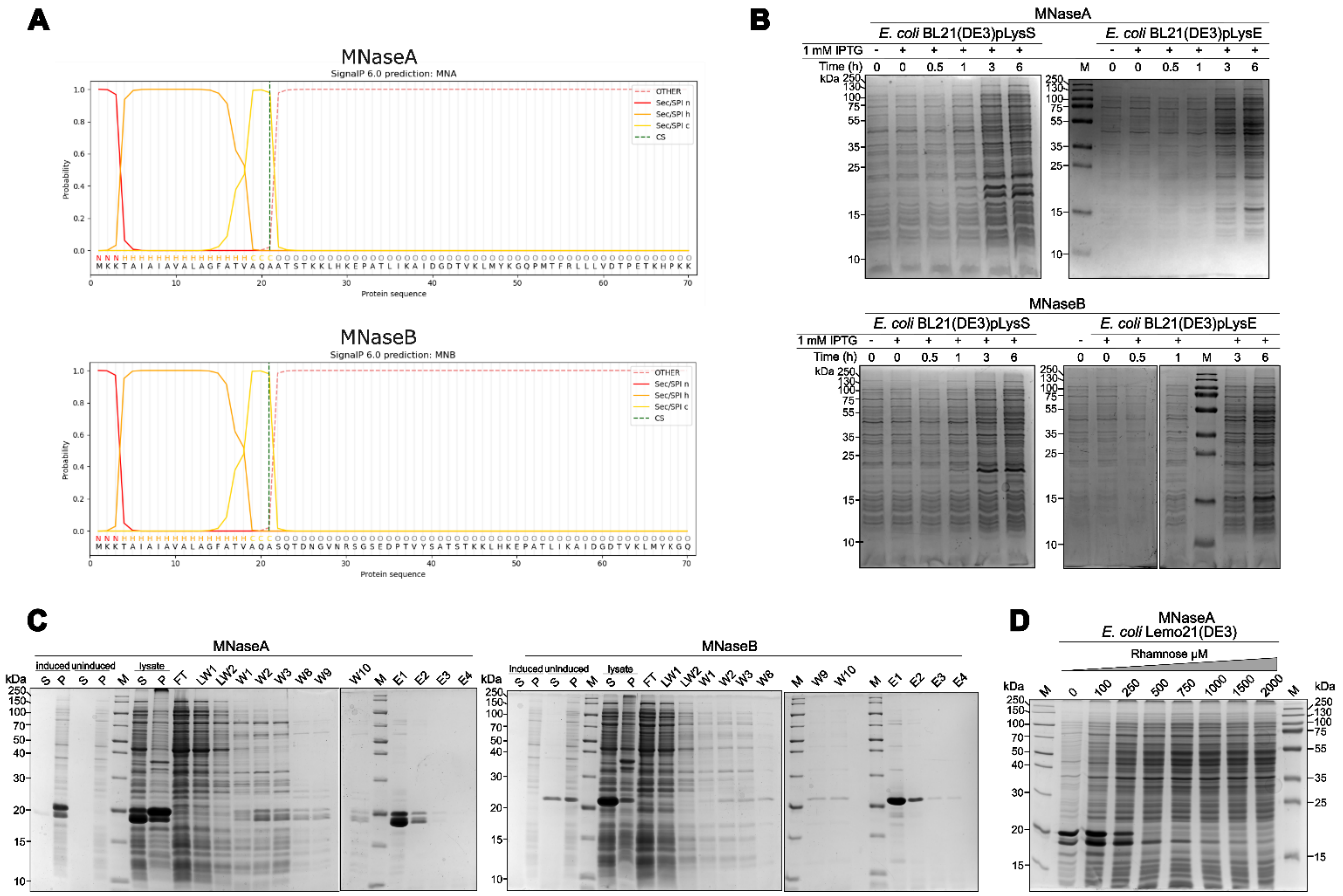


**Supporting Figure S2. (A)** Prediction of cleavage probabilities of OmpA signal peptide from MNaseA and MNaseB. Values calculated by SignalP ver 6.0. **(B)** Expression of MNaseA (top) and B (bottom) at different time points in two different *E. coli* expression strains. M – protein marker. **(C)** SDS-PAGE analysis of fractions from purification of MNaseA (left) and B (right). Induced – cultures with 1 mM IPTG, uninduced – no IPTG, S – supernatant, P – pellet, FT – flowthrough, LW – lysis buffer wash, W – wash, E – elution. **(D)** Optimization of MNaseA expression in *E. coli* Lemo21(DE3) strain by increasing rhamnose.

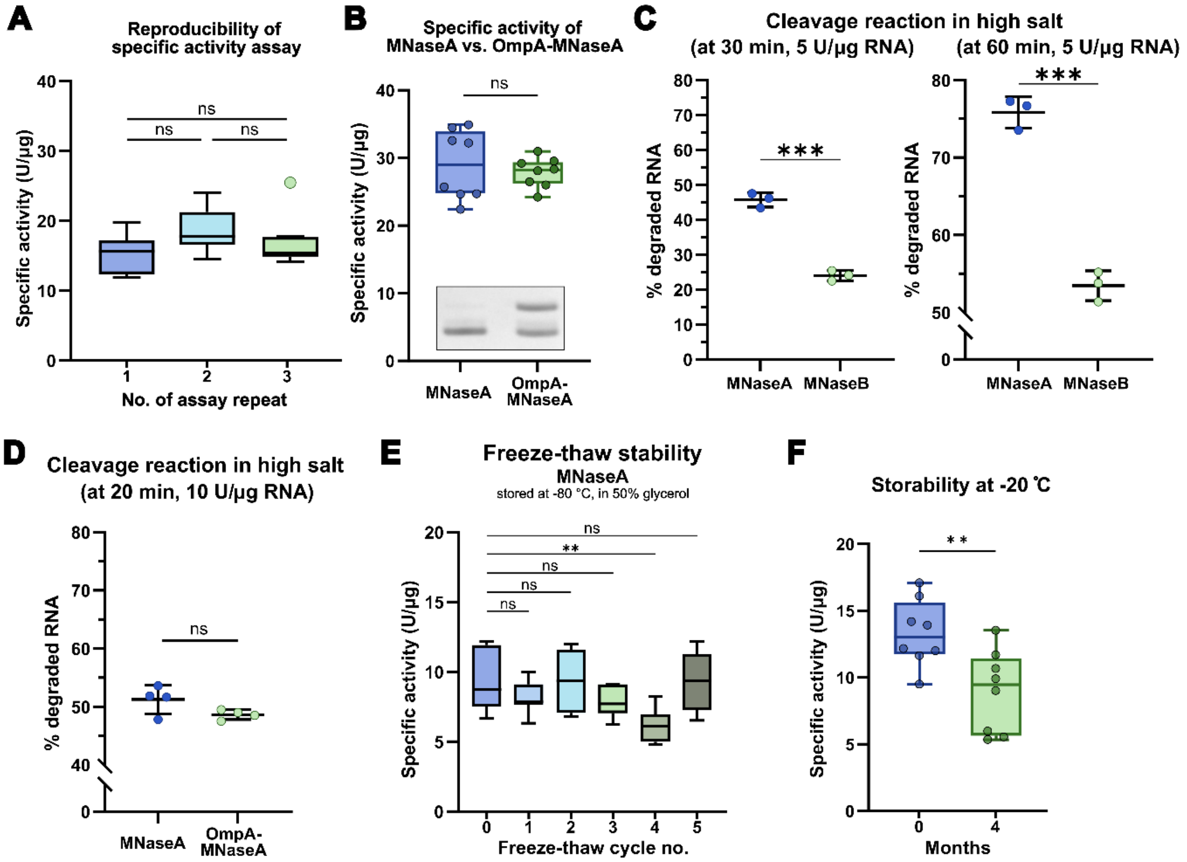


**Supporting Figure S3. (A)** Comparison of specific activity assays from three independent replicates done with commercial MNase. Ordinary one-way ANOVA (N=8, ns – not significant). **(B)** Specific activity of purified MNaseA and OmpA-MNaseA. The insert shows the purity of each tested MNaseA. Unpaired two-tailed Student's t-test (N=8, ns – not significant) **(C)** Densitometrical quantification of MNaseA and B activity in high salt reactions using 5U of enzyme per 1 µg of RNA at 30 min (left panel) and 60 min (right panel). Samples were compared by unpaired two-tailed t-test (N=3; *** p<0.001). **(D)** Densitometrical quantification of MNaseA and OmpA-MNaseA (same proteins as in panel B) activities in high salt reactions using 10U of enzyme per 1 µg of RNA after 20 min reaction. Samples were compared by unpaired two-tailed Student's t-test (N = 3; *** p<0.001). **(E)** Freeze-thaw stability of the purified MNaseA stored at -80 °C in storage buffer containing 50% glycerol. Samples were compared by ordinary one-way ANOVA (N=8, ** p=0.0039). **(F)** Stability of the purified MNaseA at -20 °C for 4 months in storage buffer containing 50% glycerol. Unpaired two-tailed Student's t-test (N=8, ** p=0.0075). Error bars in all panels represent standard deviation.

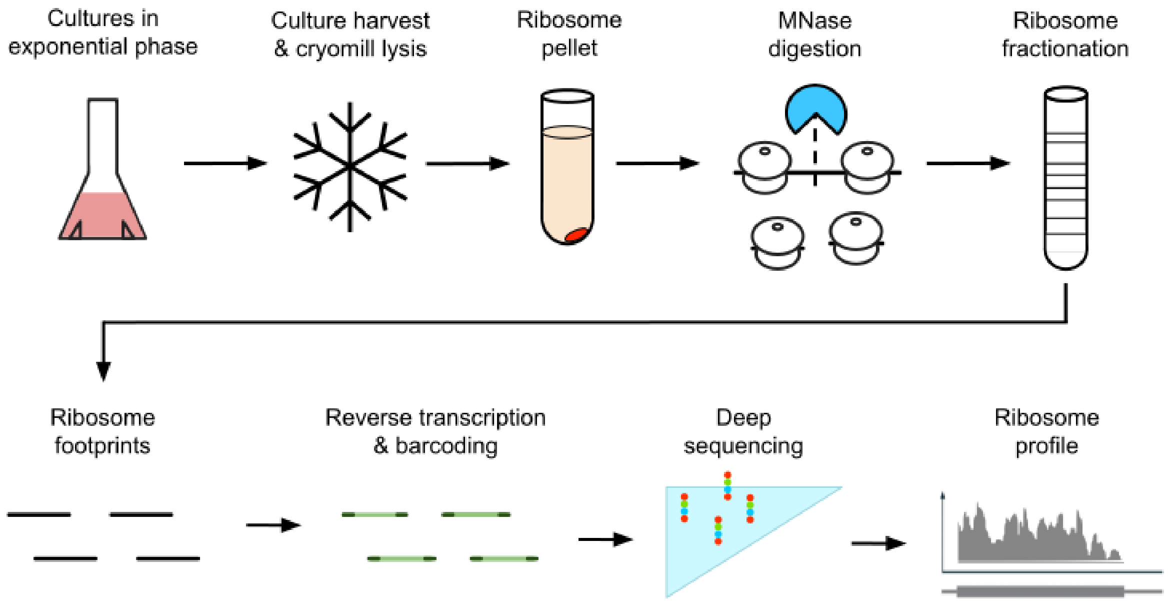


**Supporting Figure S4.** Workflow schematic for ribosome profiling in *Haloferax volcanii*. Cells were harvested in exponential phase by flash-freezing cultures dropwise and cryogenically milled (cryomilling) in liquid nitrogen. Ribosomes were pelleted from thawed lysate by ultracentrifugation, treated with MNase to digest unprotected mRNA, and separated along a 10-50% sucrose gradient; during these steps, ribosome pellets were processed in lysis buffer containing 3.4 M KCl to keep 70S monosomes intact. Ribosome footprints were then isolated from 70S sucrose fractions, reverse transcribed, barcoded with sample-specific index sequences, and sequenced.

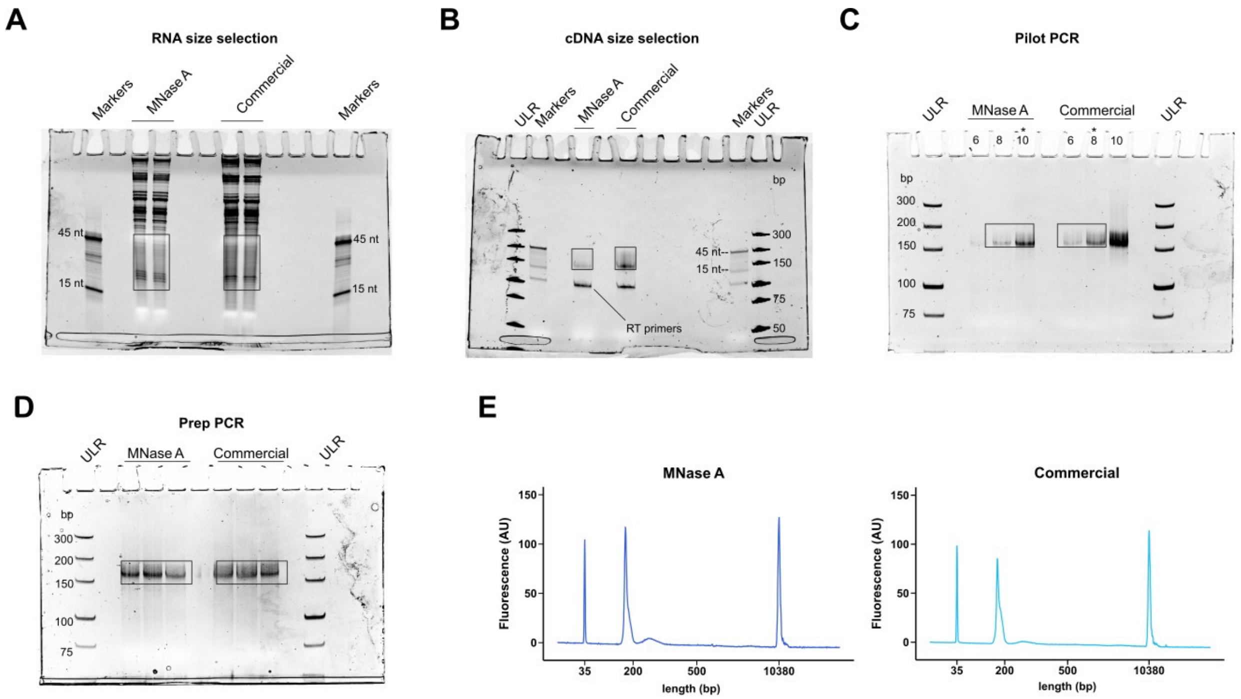


**Supporting Figure S5.** Ribosome profiling library construction steps and quality control for samples digested with MNase A or commercial MNase. Polyacrylamide gels used for **(A)** RNA footprint size selection, **(B)** barcoded/reverse-transcribed footprint size selection, **(C)** pilot PCR (cycle number selected for preparative PCR marked by (*)), and **(D)** preparative PCR. **(E)** Bioanalyzer traces of completed ribosome profiling libraries; a peak at the expected size for each final library is at ~170 bp. ULR: Ultra Low Range DNA ladder. One out of three replicates for each enzyme are shown here.

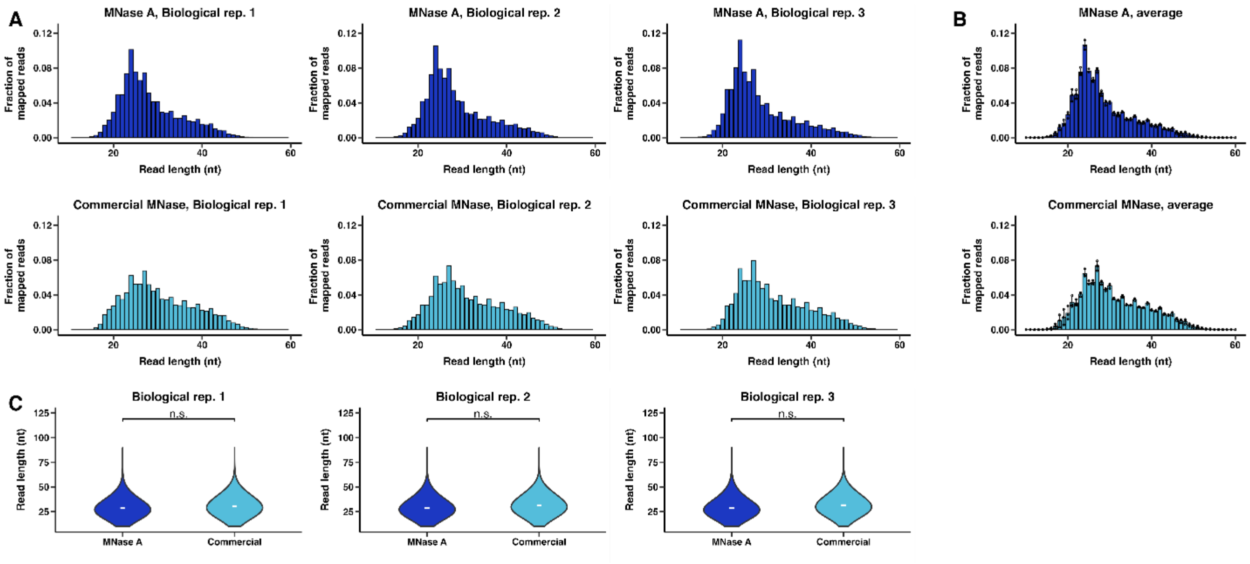


**Supporting Figure S6.** Read length distributions of ribosome profiling libraries digested with MNase A (dark blue) or commercial MNase (light blue). **(A)** Read length distributions for each of three replicates; reads per length were normalized to the total number of genome-mapped reads. **(B)** Read length distributions from MNase A and commercial MNase averaged across biological three biological replicates each; error bars indicate standard deviation. Individual biological replicates represented by points. **(C)** Read length distributions for each of the three replicates as violin plots; distribution means are marked by (-). Statistical comparison between distributions in each biological replicate (permutation test with distribution mean as test statistic, 10,000 resamples, independent) showed no significant difference (p-values: biological rep. 1, p-value = 0.7677; biological rep. 2, p-value = 0.7249, biological rep. 3, p-value = 0.6657).

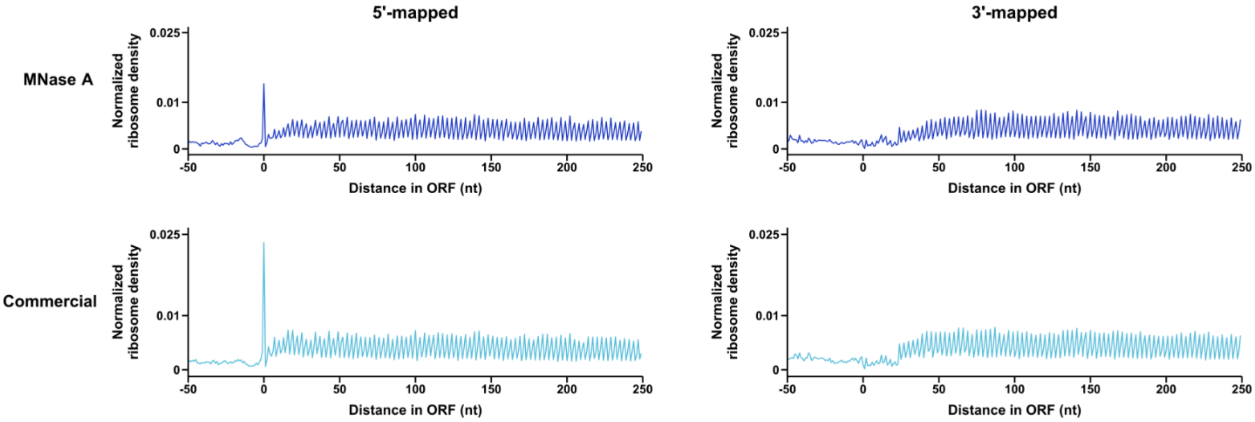


**Supporting Figure S7.** Metagene plots of ribosome profiling libraries for a single biological replicate digested with MNase A (top, dark blue) or commercial MNase (bottom, light blue). Metagene plots show the mean ribosome density on all genes aligned at their start codon using 5’-end or 3’-end mapped reads. Note that left-side panels (5’-mapped reads) are a reproduction of Figure 3D and are included here to facilitate comparison to 3’-mapped reads.

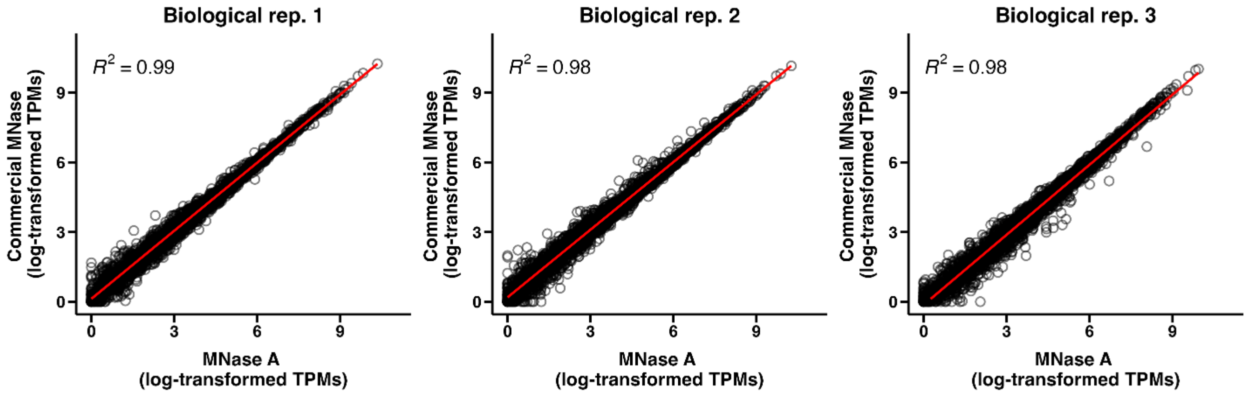


**Supporting Figure S8.** Correlation plots of ribosome occupancy per gene between samples treated with MNase A (x-axis) and commercial MNase (y-axis) within each of three biological replicates. For each gene, ribosome occupancy data from ribosome profiling were converted to transcripts per million (TPMs), log-transformed, and compared between MNase treatments.
